# Cross-domain confidence reliability and remappability of frozen single-cell representations

**DOI:** 10.64898/2026.08.31.748446

**Authors:** Guangzheng Weng, Danfei Zhu, Yufei Zhao, Patrick C Martin, Hyobin Kim, Junghyun Jung, Gi-Hoon Nam, Kyoung Jae Won

**Author notes:** These authors contributed equally to this work (Gi-Hoon Nam, Kyoung Jae Won). **Corresponding authors** Gi-Hoon Nam, E-mail address, Kyoung Jae Won.

## Abstract

Single-cell foundation models are increasingly adopted for downstream applications such as cell-type prediction. However, these predictions are often utilized without assessing their reliability, or by relying on a simple cutoff applied to the maximum softmax probability (MSP) derived from the classifier. This raises a critical question: can raw MSP be trusted as a reliable measure of confidence on unseen data (target) exhibiting diverse technical and biological variations? To address this, we introduce an audit framework to systematically evaluate the confidence estimates derived from the training set (source) against the realities of the target test set. We demonstrate that raw MSP fundamentally fails to represent true confidence when applied to target datasets. By decomposing the confidence gap between source and target data, we reveal that these discrepancies stem from a combination of rank-ordering errors and systemic probability drift. We further show that the drift can be successfully recalibrated by leveraging a small subset of labeled target data. While this recalibration improves the reliability of automated acceptance, it inherently introduces a trade-off by increasing the volume of cells requiring manual review. Ultimately, our framework establishes that the safe deployment of these models necessitates target-specific confidence adjustment using representative local labels.

## Introduction

A single-cell foundation model is a large-scale pre-trained model to learn a universal representation of cellular biology using single cell transcriptomics data. [1,2]. By learning cell embeddings from large transcriptomic datasets, these models are applied to downstream tasks such as cell-type annotation, data integration and perturbation modeling [1,3,4]. In practice, these models are typically deployed either by freezing them as feature extractors with a task-specific classifier attached, or by fully fine-tuning their parameters for the new task [5–8]. These task-specific pipelines are then applied to target cohorts containing cells from different groups of biological individuals. Throughout this study, a donors refers to one unique biological individual, regardless of how many samples or cells that individual contributes. The target cohort may differ from the source data in population composition, disease status, study site, experimental workflow or sequencing technology [4]. This practice raises a critical question: Can we reliably accept these predictions when a model is deployed on a completely new cohort?

For cell-type annotation using a frozen model, each cell in the target set receives a predicted cell type and an associated maximum softmax probability (MSP), which serves as the model’s confidence score. [9]. Typically, predictions with an MSP above a predefined cutoff are accepted automatically, while those falling below the threshold are flagged for manual review. However, a high MSP does not guarantee accuracy, nor does the raw MSP value strictly equate to the true probability of a correct prediction, especially when the model is applied to an unseen target cohort [9,10]. Furthermore, conventional benchmark evaluations use target reference labels to calculate accuracy and/or macro-F1 based on the predicted class associated with the MSP [3,4,11–13]. These metrics assess whether the predicted cell types are correct, but they fail to capture whether the model’s confidence scores are actually reliable.

Cross-domain deployment can degrade MSP reliability due to technical and biological variations. This degradation often stems from rank-ordering failures and/or confidence-mapping drift. The ordering failure happens when the model’s internal ranking system breaks down. Higher confidence scores are no longer a reliable sign that the model is right. It can happen when the new dataset introduces fundamental biological or technical shifts, such as entirely novel cell types, different disease states, or unfamiliar sequencing technologies. Conversely, confidence-mapping drift happens when the absolute confidence scores shift without changing the underlying rank-order of the target cells [10,14–16]. For example, if 90 out of 100 predictions with an MSP near 0.90 are correct in the source cohort, the mapping establishes a 90% probability of correctness. On the new data, however, that same 0.90 score might only guarantee 70% accuracy. It can happen when the new dataset experiences milder, more uniform shifts, such as changes in the relative proportions of cell types or subtle technical variations like differing sequencing depths. Because the core biological signals remain intact, the model can still correctly sort the predictions from most to least likely. If the model is just suffering from drift, reference labels from a small number of target donors can be used to re-estimate how MSP relates to prediction correctness in the target cohort. We call this process confidence recalibration. Recalibration does not change the predicted cell type or the MSP produced by the classifier; it changes the estimated probability associated with that MSP.

To systematically investigate this degradation in reliability, we audited multiple single-cell RNA sequencing (scRNA-seq) foundation models across diverse cross-domain scenarios. We partitioned datasets into distinct source (training) and target (test) cohorts. The classifier attached to the scRNA-seq foundation model is trained using the source data. We deployed frozen classifiers to target data and systematically decomposed the resulting errors into rank-ordering failures versus confidence-mapping drift. Crucially, we demonstrate that when the underlying rank-order is preserved, confidence recalibration using labeled target donors can improve the accuracy of the estimated probabilities. We further establish the minimum number of local target donors required to achieve this recalibration, and quantify how this intervention safely enables automated annotation while optimizing the downstream manual review burden. Ultimately, our framework establishes a necessary standard for evaluating and deploying reliable foundation models in single-cell transcriptomics.

## Results

### A systemic framework to audit confidence during cross-domain deployment

In each source-to-target deployment, we trained a cell-type classifier in a source cohort and applied it to a separate target cohort. We then tested whether its confidence remained reliable in the target cohort (Fig. 1A). Each released foundation model generated embeddings for source and target cells without further training. Using only cells from source-cohort donors assigned to classifier training, we standardized the embeddings and trained a multinomial logistic regression classifier (Methods). After training, no part of the annotation pipeline was updated. We used the same pipeline to predict cell types in the target cohort. Target reference labels were not used to make these predictions. For each target cell, the classifier returned a predicted cell type and its maximum softmax probability (MSP). We evaluated ten pretrained models: Geneformer, tGPT, GeneCompass, scFoundation, UCE, SCimilarity, Nicheformer, CellPLM, scGPT and scPRINT[5– 8,17–22]) and a principal component analysis (PCA) as a non-pretrained baseline [23].

**Fig. 1.**
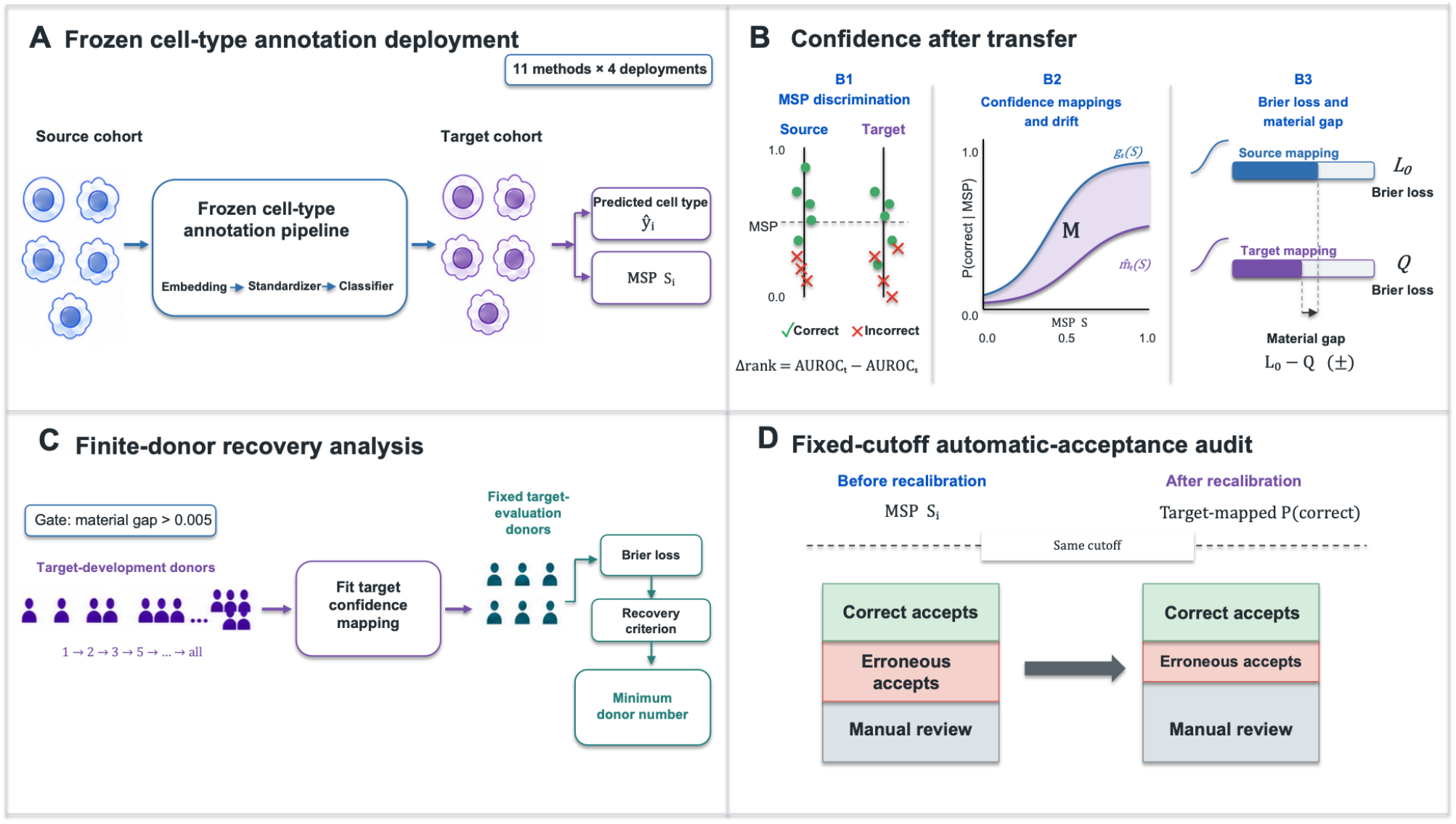
Confidence audit framework for frozen cell-type annotation pipelines. **A**, Eleven methods across four data sources makes 44 deployments. Training a classifier attached to a foundation model is done using 60% of source donors. Each frozen pipeline outputs a predicted cell type and MSP for every source and target cell. To avoid overfitting, the assigned data for evaluation is used for source and the target. **B**, The AUROC rank gap measures the change from source to target in the ability of MSP to discriminate correct from incorrect predictions (B1). *M* measures the difference between the source and target confidence mappings (B2). *L*_0_ is the Brier loss under the source confidence mapping, whereas *Q* is the Brier loss under the target confidence mapping. Both are evaluated on the same target cells. Their difference, *L*_0_ − *Q*,is the reduction in Brier loss defined as the material gap (B3). **C**, Deployments with a material gap > 0.005 undergo finite-donor recovery analysis to estimate the minimum donor number. **D**, Automatic acceptances, erroneous automatic acceptances and manual review are compared before and after recalibration using the same cutoff.

These were tested using four resources: the Asian Immune Diversity Atlas (AIDA), the Kidney Precision Medicine Project (Kidney), the Human Lung Cell Atlas (HLCA) and published retinal studies[24–28]; the retinal analysis additionally used raw count matrices from GEO accession GSE158629[29]. To systematically capture distinct biological and technical domain shifts, we partitioned these datasets into specific source (training) and target (test) cohorts. Specifically, we evaluated demographic transfer using AIDA (training on a South Korean cohort and applying to a Thai target cohort[24]), disease-state transfer in the Kidney dataset (normal kidney to chronic kidney disease [30,31]) and HLCA[26] (healthy to interstitial lung disease), and technological transfer in the Retina dataset (healthy scRNA-seq to single nucleus RNA sequencing (snRNA-seq) data [27,28]). For each source dataset, we used 60% of them to train the classifier while the remaining 40% were used for independent recalibration (20%) and evaluation (20%) to avoid overfitting and information leakage. Similarly, we only used 30% of the target datasets for evaluation of the performance and remaining 70% for recalibration.

We first assessed the ability of MSP to discriminate between correct and incorrect predictions in both the source and the target cohort (Fig. 1B1). For this assessment, we used the evaluation set from donor and the target. For each deployment, we calculate the area under the receiver operating characteristic curve (AUROC) using the MSP score and a binary outcome for prediction correctness (*C*_*i*_) (*C*_*i*_ = 1 if matches and *C*_*i*_ = 0 otherwise). Then, there is an AUROC rank gap between the target and the source is

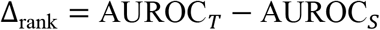

We utilized the AUROC rank gap to quantify the degradation in the model’s ability to distinguish correct from incorrect predictions following the cross-domain deployment.

The confidence mapping denotes the mathematical function that converts a specific MSP score into the true probability of a correct prediction. Both the source and target mappings are fitted independently via a natural-cubic spline estimator [15,16] (Methods). For this mapping, we used the calibration sets reserved from the source (20%) and the target (70%). After both mappings are fixed, we apply them to cells from target donors used only for evaluation. For each cell *i*, the same MSP *S*_*i*_ is passed through the source and target confidence mappings to obtain *g*_*s*_(*S*_*i*_) and 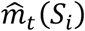, respectively (Fig. 1B2). We define confidence-mapping drift, M, as the weighted mean of the squared differences between these two estimates.

To evaluate the two confidence mappings, we apply both mappings to the same cells from target donors used only for evaluation. For each cell *i* with MSP *S*_*i*_, the source confidence mapping gives *g*_*s*_(*S*_*i*_), whereas the target confidence mapping gives 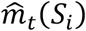. The reference label determines whether the predicted cell type is correct (*C*_*i*_ = 1) or incorrect (*C*_*i*_ = 0). For each mapping, we calculate the squared difference between its estimated probability and *C*_*i*_ for every cell. We first averaged these squared errors within each donor and then averaged the donor-level values equally across donor (Methods). The resulting donor-equal Brier loss is denoted by *L*_0_ under the source confidence mapping and by *Q* under the target confidence mapping[32]. Thus, *Q* quantifies how closely the probabilities reported by the target confidence mapping agree with the observed 0/1 correctness outcomes. *M* measures the difference between *g*_*s*_(*S*_*i*_) and 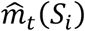 across the same cells (Methods). These quantities satisfy

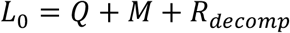

where *R*_decomp_ is the signed cross-term produced by the squared expansion (Methods). We define the reduction in Brier loss obtained by replacing the source confidence mapping with the target confidence mapping as the material gap:

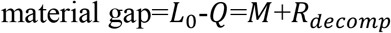

We used the material gap to decide which method–deployment combinations to study further (Fig. 1C,D). A combination entered the finite-donor recovery analysis when its material gap was strictly greater than 0.005. This analysis asked how many target donors were needed to reproduce the reduction in Brier loss obtained using all target donors available for recalibration. For each combination that passed this threshold, we selected increasing numbers of target donors and used the reference cell-type labels for their cells to refit the target confidence mapping. We tested each fitted mapping on cells from a separate group of target donors used only for evaluation (Methods). We recorded the smallest tested donor count at which the recovery criterion was met and remained met at all larger tested counts. Finally, at fixed cutoffs, we compared direct use of MSP with recalibrated probabilities and counted erroneous automatic acceptances and cells sent for manual review (Fig. 1D).

### Prediction accuracy after cross-domain deployment does not establish the reliability

We initially evaluated MSP’s performance by measuring the rate of erroneous automatic acceptances per 1,000 cells across the 44 target deployment scenarios. Overall, we observed high average MSPs with 35 of the 44 deployments maintaining a cell-type prediction accuracy of 0.80 or greater. However, contrary to the expectation that a higher MSP correlates with reduced error, we found that increased cell-type prediction accuracy is not consistently accompanied by a decrease in erroneous automatic acceptances (Spearman *ρ* = 0.08 ) (Fig 2A). For example, although Geneformer achieved similar accuracies for the AIDA and HLCA cohorts (0.880 and 0.884, respectively), but the corresponding numbers of erroneously accepted predictions per 1,000 cells differed drastically (10.1 for AIDA versus 38.3 for HLCA) and their overall automatic-acceptance proportions also diverged significantly (62.2% and 83.0%, respectively). Furthermore, even within the same HLCA dataset, SCimilarity and Geneformer achieved comparable prediction accuracies (0.886 and 0.884, respectively) and overall automatic-acceptance proportions (82.6% and 83.0%). This demonstrates that methods with seemingly equivalent accuracy and acceptance rates can still exhibit drastically different rates of confident errors. Consequently, standard benchmarks relying solely on overall prediction accuracy overlook the true volume of erroneous predictions that clear the MSP cutoff.

**Fig. 2.**
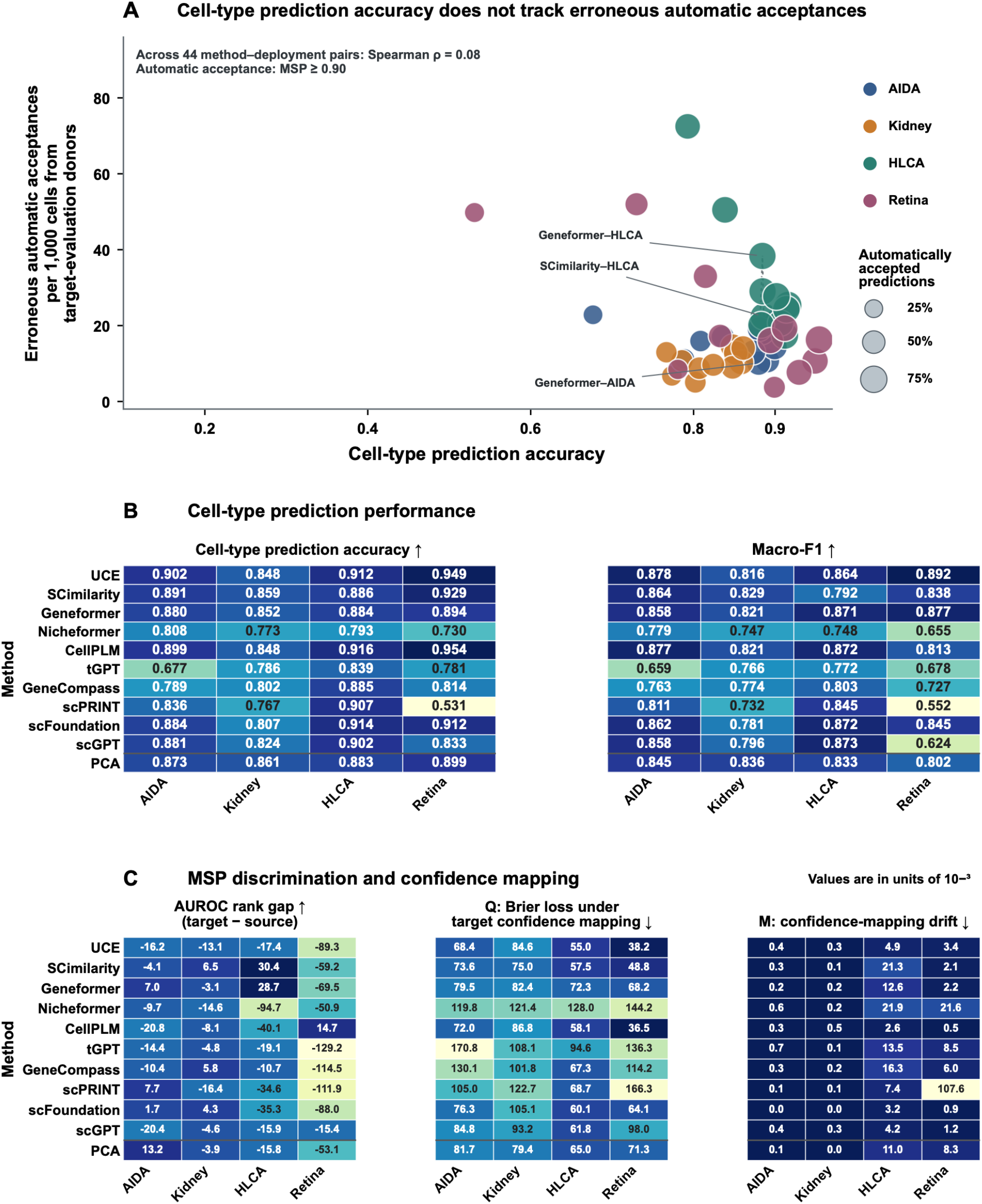
Cell-type prediction performance does not capture MSP reliability after cross-cohort deployment. **A**, Each bubble represents one method–deployment combination. The x-axis shows donor-equal cell-type prediction accuracy. The y-axis shows erroneous automatic acceptances per 1,000 cells when predictions with MSP ≥ 0.90 are accepted automatically. Bubble area represents the proportion of predictions accepted automatically. Each metric is first calculated within each target donor used only for evaluation and then averaged equally across donors. Spearman *ρ* is calculated across all 44 method–deployment combinations. **B**, Cell-type prediction accuracy and macro-F1 for 11 methods across four deployments. Accuracy uses donor-equal weights. Macro-F1 is the unweighted mean of class-specific F1 values across the complete cell-type set. Higher values indicate better cell-type prediction performance. **C**, AUROC rank gap, *Q* and *M* across the same 44 combinations. The AUROC rank gap is target AUROC minus source AUROC; positive values indicate improved MSP discrimination in the target cohort, whereas negative values indicate reduced discrimination. *Q* is the donor-weighted Brier loss under the target confidence mapping, calculated using cells from target donors used only for evaluation. *M* measures the donor-weighted difference between the probabilities reported by the source and target confidence mappings for the same cells. Lower *Q* indicates smaller probability error, whereas lower *M* indicates less confidence-mapping drift. Values in C are reported in units of 10^−3^.

While deployments on the AIDA, Kidney, and HLCA datasets achieved high accuracies near or above 0.80, performance on the Retina cohort, which is to deploy across technologies (scRNA-seq to snRNA-seq), diverged significantly (ranging from 0.531 to 0.954). To evaluate the degradation of model confidence, we further investigated the AUROC rank gap, *Q* (the baseline target error), and *M* (the confidence-mapping drift) across the deployments (Fig. 2C). For AIDA and Kidney dataset, the source and target confidence mappings were similar (*M*< 0.7 × 10^−3^) but with a divergent *Q* (ranging from 68.4 × 10^−3^ to 170.8 × 10^−3^). Furthermore, the AUROC rank gap, which measures the ability to discriminate between correct and incorrect predictions, exhibited both positive and negative values. These distinct patterns highlight that cell-type prediction accuracy, AUROC rank gap, *Q*, and *M* measure independent phenomena.

Despite the complexity of these foundation models, they do not consistently exceed the cell-type prediction accuracy of a simple PCA baseline (Fig. 2B; Supplementary Fig. 1A). Intriguingly, model deployments with performance inferior to PCA demonstrated higher Q values (ρ = 0.95). This tells that the foundation models cannot perform better than PCA if the deviation of data is large enough to change the confidence rank order. Compared with retaining 128 principal components, retaining 64 improves accuracy and reduces *Q* in all four deployments, whereas its effect on *M* is inconsistent. No single PCA dimensionality performs best across all deployments and metrics (Supplementary Fig. 1B).

### Evaluating the Material Gap and Minimum Donor Requirements for Recalibration

To evaluate practical benefits of recalibration, we quantified improvement (or material gap) for all deployment. Among 44 deployments, fourteen (ten from HLCA and four from Retina) showed a material gap greater than the 0.005 threshold (Fig. 3A). In the scPRINT–HLCA deployment, the Brier loss for the source (*L*_0_ =0.0797) and the target (*Q* = 0.0687) showed a material gap of 11.1 × 10^−3^. scPRINT–Retina deployment showed the greatest material gap of 117.6 × 10^−3^ (Fig 3A and Supplementary Table 1). scPRINT–Retina deployment still retains the highest *Q* (0.1663) among the 14 pairs after recalibration. This contrast demonstrates that a single foundation model can achieve substantially different reductions in error depending on the target dataset, and that a massive relative improvement does not guarantee a low absolute error.

**Fig. 3.**
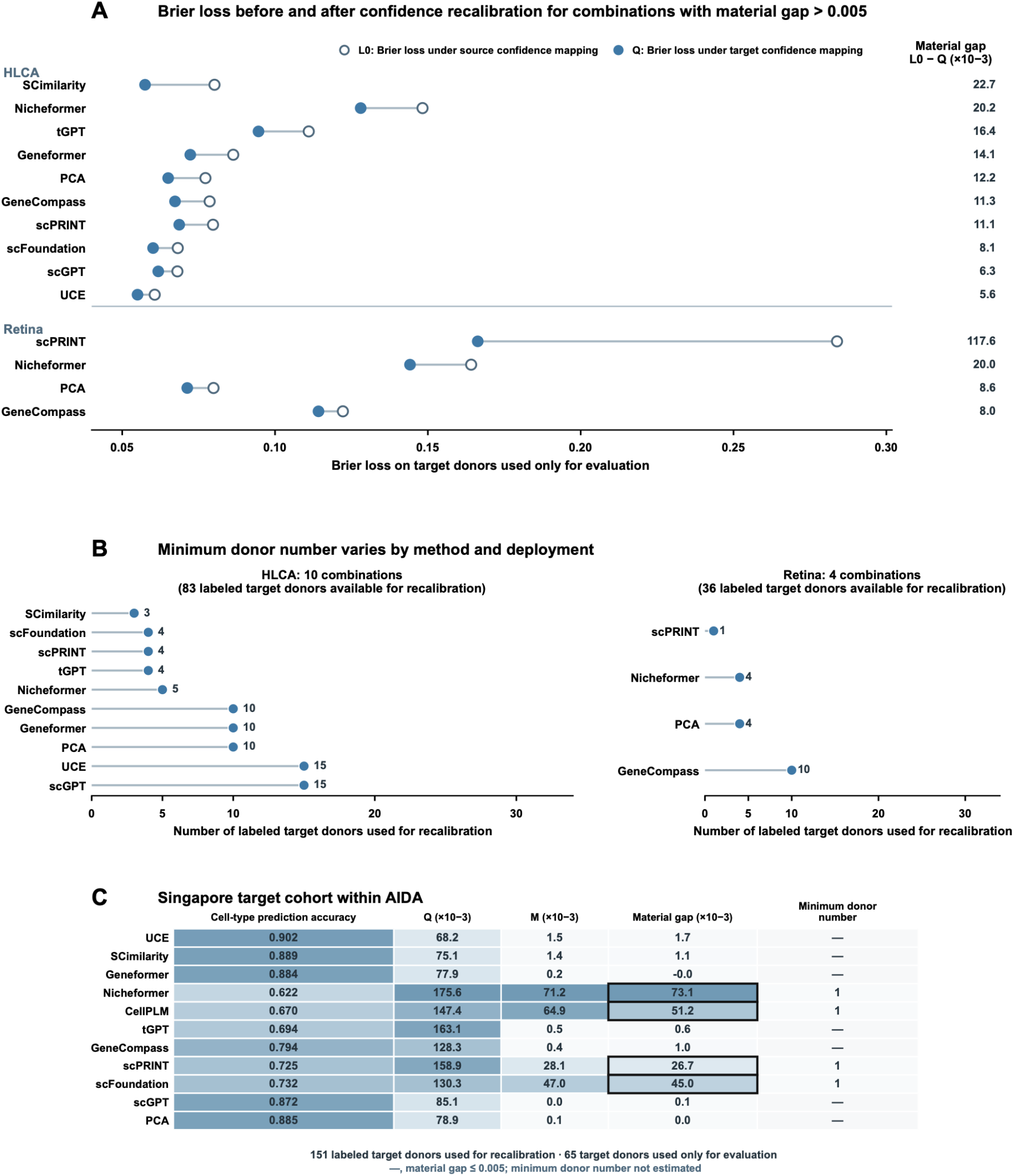
Changes in Brier loss after confidence recalibration and finite-donor recovery analysis. **A**, *L*_0_, *Q*, and the material gap (>0.005) for the 14 deployment. Open circles indicate *L*_0_, whereas filled circles indicate the Brier loss produced by the target confidence mapping (*Q*). **B**, Minimum donor number for improvement for the 14 pairs above cutoff. **C**, Evaluation of the Singapore target cohort using the same AIDA source cohort and frozen cell-type annotation pipelines. Columns report cell-type prediction accuracy, *Q, M*, material gap, and minimum donor number, respectively. Black outlines identify methods with a material gap greater than 0.005. A dash indicates that the method does not enter the finite-donor recovery analysis.

We further investigated the number of donors needed to reduce the material gap. For each deployment above a cutoff, we fit the target confidence mapping with a natural-cubic spline while increasing the number of target donors with label (Method). A donor count met the recovery criterion only when repeated fits consistently reduced Brier loss below *L*_0_ and recovered a prespecified proportion of the reduction obtained using all available labeled target donors.

Among the donor counts tested, we defined the first count that met the recovery criterion (Method) and continued to meet it at all larger tested counts as the minimum donor number. Eight of the 14 deployment with a material gap >0.005 required no more than five labeled target donors (Fig. 3A,B). To understand the broader applicability of our recalibration approach, we next examined whether the benefits of target confidence mapping depended on the target population. To assess the impact of the target population, we evaluated independent target cohorts by using Singapore cohorts from the AIDA dataset while keeping the South Korean cohorts as the training set (Fig. 3C).

Compared with previous deployment to Thai cohorts where no deployment passed the cutoff due to low material gap, Nicheformer, CellPLM, scPRINT, and scFoundation met this criterion for the deployment to Singapore donors (>0.005). Notably, when using a natural-cubic spline, all four successful methods reached the minimum donor number of 1 to fix the model’s confidence using only one donor (Fig. 3C). These results show that, for methods with a material gap above the threshold in Singapore, recalibration using a single labeled target donor was sufficient to consistently reduce Brier loss on target donors used only for evaluation.

### Manual-review trade-offs after confidence recalibration

While recalibration effectively limits false acceptances, it can simultaneously drive a higher volume of predictions below the automated cutoff, thereby escalating the demand for manual review. To quantify this trade-off, we investigated two key questions: First, does confidence recalibration demonstrably reduce the incidence of automated misclassifications? Second, what is the magnitude of additional manual intervention required to achieve this performance gain?

We first examined whether method–deployment combinations with a material gap greater than 0.005 also produced more erroneous automatic acceptances when MSP was used directly (Fig. 4A). We divided the combinations into two groups according to whether their material gap was greater than 0.005 or at most 0.005. For each group, we calculated erroneous automatic acceptances per 1,000 cells at MSP cutoffs of 0.80, 0.90 and 0.95. Increasing the MSP cutoff reduced erroneous automatic acceptances in both groups because fewer predictions were accepted automatically. At every MSP cutoff, however, combinations with a material gap greater than 0.005 had more erroneous automatic acceptances than those with a material gap at or below 0.005. The same pattern was observed when the South Korea-trained pipelines were applied to the Singapore target cohort (Fig. 4A).

**Fig. 4.**
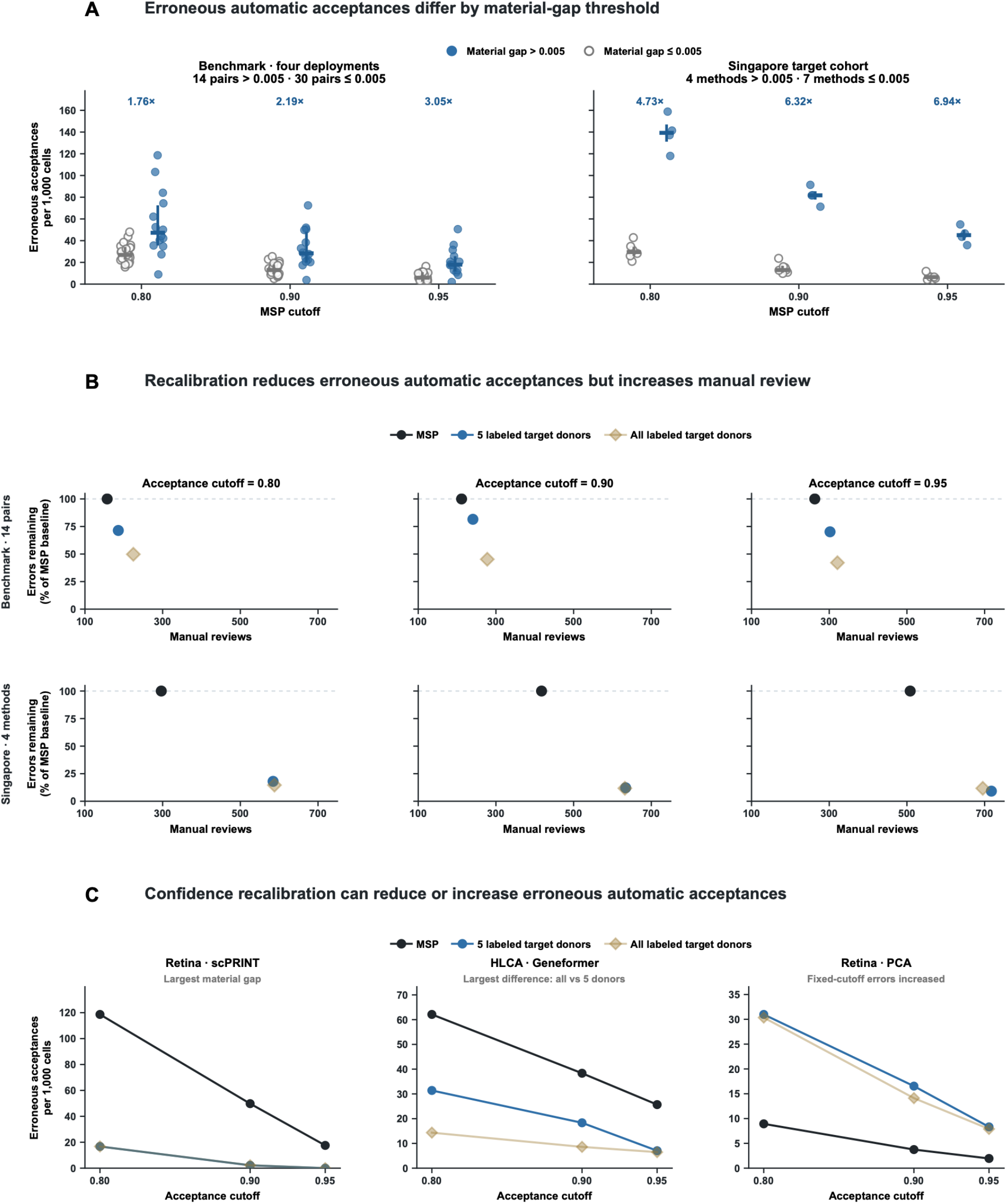
Decision consequences of confidence recalibration at fixed cutoffs. **A**, Erroneous automatic acceptances when MSP was used directly as the decision score. Groups were defined by material gap > 0.005 or ≤ 0.005. Each point represents a benchmark model–deployment combination or a Singapore method. Horizontal lines show medians, and vertical lines show interquartile ranges. Annotated values give the ratio between the two group medians. Cutoffs were 0.80, 0.90 and 0.95. **B**, Trade-off between erroneous automatic acceptances and manual review. At each cutoff, erroneous automatic acceptances under MSP were set to 100%; recalibrated results show the percentage of these errors that remained. Manual review is reported per 1,000 cells from target donors used only for evaluation. Markers show medians across the 14 benchmark combinations or four Singapore methods with material gap > 0.005. Target confidence mappings were fitted using a natural-cubic spline and either five or all available labeled target donors. Five-donor results were summarized across 1,000 donor samples. **C**, Three examples showing reduced erroneous automatic acceptances differences between recalibration using five and all labeled target donors, increased erroneous automatic acceptances, and high erroneous automatic acceptances when material gap was ≤ 0.005. Each subpanel uses an independent vertical-axis range. Before recalibration, MSP was used as the decision score. After recalibration, the estimated probability of correctness from the target confidence mapping was used instead. The same numerical cutoff was applied in both cases. Predicted cell types and MSP values remained unchanged.

We next examined how confidence recalibration and the number of target donors used to fit the mapping changed erroneous automatic acceptances and manual review (Fig. 4B). We compared direct use of MSP with two recalibrated decision rules. One mapping was fitted using cells from five target donors. The other used cells from all target donors available for recalibration. Both mappings were fitted from the cells’ MSP values and whether the predicted cell type matched the reference label. Each rule was evaluated on cells from separate target donors used only for evaluation. For each cutoff, we set the number of erroneous automatic acceptances under MSP to 100%. Errors remaining after recalibration were expressed relative to this baseline, while manual reviews were reported per 1,000 cells. Across the 14 benchmark method–deployment combinations with a material gap greater than 0.005, both recalibrated rules reduced the median percentage of errors remaining but increased manual review (Fig. 4B). In the benchmark, the mapping fitted with all available target donors reduced errors more than the mapping fitted with five donors. In the South Korea-to-Singapore deployment, both mappings reduced errors relative to MSP, but using all available target donors did not consistently improve on using five donors.

The Retina–scPRINT deployment, which exhibited the largest material gap, showed a marked reduction in erroneous automatic acceptances as the confidence cutoff increased, and recalibration substantially improved these outcomes(Fig. 4C). The HLCA–Geneformer deployment demonstrated a similar, though less dramatic, reduction in the error rate. In contrast, recalibration of the Retina-PCA deployment actually increased the error rate after recalibration, indicating that this recalibration approach is primarily effective for foundation models.

In sum, implementing confidence-based automated annotation requires the trade-off between error reduction and increased manual review. Furthermore, confidence recalibration cannot rescue a fundamentally poor classifier, highlighting the necessity of robust baseline predictions prior to deployment.

## Discussion

Single-cell foundation models are increasingly deployed across diverse biological and technical domains. However, they are frequently utilized without rigorous consideration of their predictive confidence. While these models acquire extensive prior knowledge through large-scale pretraining, they are often applied to unseen target datasets under the implicit assumption that their raw confidence scores remain reliable. Our study demonstrates that relying on these raw scores without a dedicated confidence audit is fundamentally flawed (Fig 2A). We find that high cell-type prediction accuracy on a target dataset does not necessarily correspond to fewer erroneous automatic acceptances at a fixed MSP cutoff (Fig 2A). Consequently, an evaluation benchmark based solely on accuracy obscures critical underlying vulnerabilities, leaving high-throughput workflows highly susceptible to the silent acceptance of overconfident, incorrect predictions.

To address this critical methodological gap, we implemented a comprehensive confidence audit framework designed to evaluate potential issues in cross-domain model deployment. Rather than treating MSPs as confidence, our framework explicitly decomposes performance by evaluating confidence-mapping drift (*M*), the Brier loss in targets (*Q*), and the operational material gap. We systematically applied this audit system to evaluate multiple single-cell foundation models across a wide array of cross-domain deployment scenarios. By assembling 44 distinct method– deployment combinations encompassing cross-cohort and cross-modality shifts (e.g., AIDA, Kidney, HLCA, and Retina), we were able to rigorously test these models. Notably, our results reveal that confidence degradation is highly deployment-specific. Furthermore, despite their massive scale, foundation models do not consistently outperform simpler baseline methods, such as PCA, across all deployments and confidence metrics, highlighting the necessity of contextual evaluation.

Through this audit, we demonstrate that when a model’s internal rank-ordering is preserved, confidence-mapping drift can be successfully recalibrated. For the 14 deployments that met our prespecified threshold criterion, replacing the source confidence mapping with an empirical target confidence mapping reduced the material gap. Intriguingly, our recovery analysis reveals that this recalibration is highly data-efficient (Fig 3,4). For several deployments, labeled data from a minimal number of target donors (often as few as one to five) was sufficient to consistently reduce prediction error on independent target cohorts.

However, our framework also highlights the operational trade-offs inherent to model recalibration. While it re-estimates the probability of correctness associated with an MSP, it does not change the underlying predicted cell type. Consequently, while recalibration safely moves many incorrect predictions below the automatic-acceptance cutoff, it may also inadvertently refer correct predictions for manual review. The operational trade-off between the reduction of erroneous automatic acceptances and the increased burden of manual review must be explicitly quantified and managed. If a model completely loses its ability to discriminate between correct and incorrect predictions, confidence recalibration alone cannot resolve the issue, and the underlying classifier must be updated.

We note specific limitations to our current findings. Our empirical evaluations are based on a restricted set of source-to-target deployments and focus specifically on frozen cell-type annotation pipelines using MSP as the sole confidence metric. The findings from this study cannot be directly extrapolated to other workflows, such as fine-tuning or nearest-neighbour label transfer[4,5,18].

By bridging the gap between raw model outputs and verifiable diagnostic reliability, our framework establishes an indispensable standard for the safe translation of single-cell foundation models into real-world applications.

## Methods

### Study design, cohorts and donor-disjoint roles

This study evaluates ten pretrained single-cell foundation models and PCA across four benchmark source-to-target deployments: AIDA, Kidney, HLCA and Retina[24,25,27,28,31] (Supplementary Methods Table M1). We evaluated 11 methods in each of four source-to-target deployments, yielding 4 method–deployment combinations. All data splits were performed at the donor level rather than the cell level. Each donor represented one biological individual, and all cells from that donor were assigned to the same role. Within each deployment, source donors were divided approximately 6:2:2 into source-training, source-calibration and source-evaluation groups. Target donors were divided approximately 7:3 into target-development and target-evaluation groups. Exact donor and cell counts for each role are reported in Supplementary Table M1. Source-training donors are used to fit the standardizer and train the classifier. Source-calibration donors are used to fit the source confidence mapping, whereas source-evaluation donors are used to evaluate source cell-type prediction and MSP discrimination. Target-development donors are used to fit the target confidence mapping and provide subsets for finite-donor recovery analysis. Target-evaluation donors are excluded from all fitting and used to evaluate target cell-type prediction, MSP discrimination, Brier loss and fixed-cutoff outcomes.

Finite-donor subsets do not constitute an additional donor role.

### Cell embeddings and frozen cell-type annotation pipelines

For each of the ten pretrained models, the same frozen parameters and its corresponding preprocessing workflow are used to generate fixed-dimensional embeddings for cells from the source and target cohorts. The detailed parameter checkpoint, preprocessing procedure, embedding extraction method and dimensionality for each method are in Supplementary Table M2.

For each deployment, source-training donors are used to fit a feature-wise standardizer and an L2-regularized multinomial logistic-regression classifier. The inverse regularization strength is selected from *C* ∈ {0.01, 0.1, 1, 10} using three donor-disjoint folds within the source-training donors. A standardizer is fitted independently within each inner fold. Model selection uses the equally weighted mean negative log probability across donor-by-true-class groups, with the smaller *C* selected in the event of a numerical tie. After *C* is selected, the standardizer and classifier are refitted using all source-training donors and then fixed.

After *C* is selected, the standardizer and classifier are refitted using all source-training donors. They are then fixed and applied without further training to all cells from donors not used for classifier training. Reference labels for these cells are not used to generate class probabilities, predicted cell types or MSP values. For cell *i*, the classifier outputs a probability *p*_*ik*_ for each candidate cell type *k*, and we define:

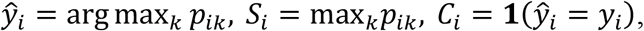

where *ŷ*_*i*_ is the predicted cell type, *S*_*i*_ is the MSP and *y*_*i*_ is the reference label. *C*_*i*_ equals 1 when *ŷ*_*i*_ = *y*_*i*_ and 0 otherwise. Reference labels are used only after prediction to determine *C*_*i*_. Subsequent analyses use *ŷ*_*i*_ and *y*_*i*_ to evaluate cell-type prediction performance, use *S*_*i*_ as the score and *C*_*i*_ as the binary outcome to evaluate the ability of MSP to discriminate between correct and incorrect predictions, and use *S*_*i*_ to estimate the probability of correctness. Class probabilities, predicted cell types and MSP values remain fixed throughout all subsequent analyses.

### Donor-weighted cell-type prediction and MSP discrimination

All evaluations except macro-F1 use donor-equal weights. For cell *i* in evaluation role *r*, let *d*(*i*) denote its donor, *π*_*i*_ its sampling inclusion probability and *D*_*r*_ the number of donors in that role. The cell weight is defined as:

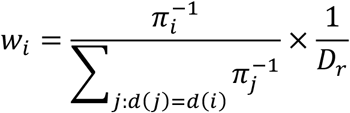

Each donor therefore has a total weight of 1/*D*_*r*_, and the weights sum to 1 within each role. For cells not subject to sampling, *π*_*i*_ = 1. These weights retain inverse-inclusion correction within each donor[34] while preventing donors with more cells from receiving greater total weight.

For evaluation role *r*, cell-type prediction accuracy is defined as:

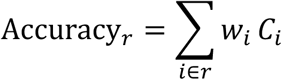

The benchmark analyses use accuracy among target-evaluation donors to assess target-domain cell-type prediction performance. Macro-F1 is defined as the unweighted arithmetic mean of the class-specific F1 values across the complete frozen class set and does not use donor-equal weights. If a class is absent from the evaluation cells or is never predicted, its F1 is set to 0.

Donor-weighted AUROC is calculated separately among source-evaluation and target-evaluation donors[35], using MSP *S*_*i*_ as the score and correctness *C*_*i*_ as the binary outcome. For evaluation role *r*:

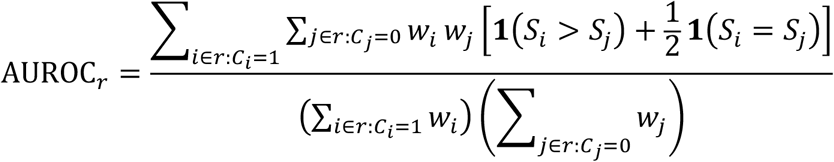

The AUROC rank gap is defined as:

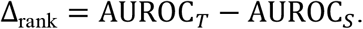

To evaluate outcomes when MSP is used directly for automatic acceptance, predictions with MSP of at least 0.90 are defined as automatically accepted. Let:

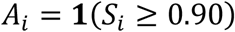

The automatic-acceptance proportion among target-evaluation donors is:

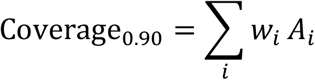

The number of erroneous automatic acceptances per 1,000 target-evaluation cells is:

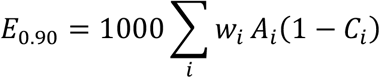

The standardized denominator is all target-evaluation cells, rather than only the automatically accepted cells.

### Confidence mappings, Brier-loss decomposition and material gap

A confidence mapping converts the MSP into an estimated probability of the cell-type prediction being accurate. The source confidence mapping *g*_*s*_ is fitted using source-calibration donors, whereas the target confidence mapping 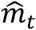 is fitted using all target-development donors:

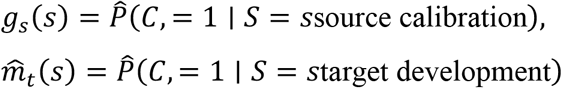

Both mappings are fitted independently using logistic regression with MSP as the sole input and a natural-cubic spline basis[15,16]. Prediction correctness *C*_*i*_ is the binary outcome, and MSP *S*_*i*_ is the only input. This spline allows the estimated probability of correctness to change smoothly and nonlinearly across the MSP range. Each donor contributes the same total weight during fitting. Knot positions are determined separately for each mapping from the MSP values in its fitting data. Three internal knots are placed at the weighted 25th, 50th and 75th percentiles. Boundary knots are placed at the weighted 1st and 99th percentiles. The source and target confidence mappings therefore have their own knots and coefficients. Before fitting, the spline-derived variables are centered and scaled using the same donor weights. The intercept is not penalized. The remaining coefficients use an *L*_2_ penalty of *λ* = 1, adjusted for the effective sample size of the weighted data. Coefficients are estimated using L-BFGS-B, with at most 1,000 iterations and a convergence tolerance of 10^−12^. Estimated probabilities are restricted to the interval [10^−6^, 1 − 10^−6^].

After both mappings are fixed, the same MSP from each target-evaluation cell is passed through both mappings. All sums below are taken over target-evaluation cells using the weights *w*_*i*_ defined above. The Brier loss produced by the target confidence mapping is defined as[32]:

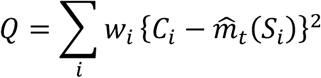

*Q* measures the overall discrepancy between the estimated probabilities of correctness produced by the target confidence mapping and the observed 0/1 correctness outcomes. A higher *Q* indicates a larger discrepancy. For example, when correct and incorrect predictions are concentrated within similar MSP ranges, the mapping can assign them only similar probabilities of correctness, leaving a high Brier loss. *Q* is an empirical loss evaluated on independent target-evaluation donors and is not a theoretical lower bound.

The difference between the source and target confidence mappings is quantified as:

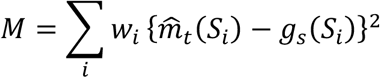

For each target-evaluation cell, both mappings receive the same MSP. *M* is the donor-weighted mean squared difference between the probabilities of correctness produced by the two mappings. A higher *M* indicates greater confidence-mapping drift. Because *M* does not use the observed correctness *C*_*i*_, it does not quantify the actual reduction in Brier loss obtained by adopting the target confidence mapping.

The Brier loss obtained by applying the source confidence mapping directly to the same target-evaluation cells is defined as:

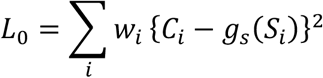

For each cell, the error produced by the source confidence mapping can be written as:

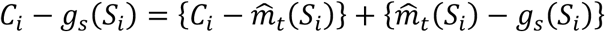

Squaring this expression and summing with donor-equal weights gives:

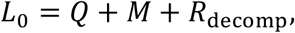

where the cross-term is:

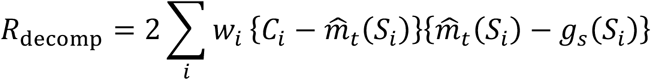

The material gap is defined as the Brier loss under the source confidence mapping minus that under the target confidence mapping:

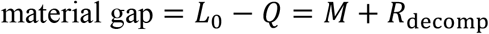

A positive material gap indicates that adopting the target confidence mapping reduces Brier loss among target-evaluation donors. A larger positive value indicates a greater reduction. *M* quantifies how much the two mappings differ, whereas the material gap quantifies the actual reduction in Brier loss after replacing the source mapping. Because *R*_decomp_ can be positive or negative, a large *M* does not guarantee a positive material gap.

### Finite-donor recovery

Only method–deployment combinations with a material gap > 0.005 under the natural-cubic spline entered the finite-donor recovery analysis.

For each included combination, we generated 1,000 reproducible random orderings of the target-development donors. At each tested donor count *k*, the target confidence mapping was fitted using the first *k* donors in each ordering and evaluated on the same target-evaluation donors. All methods within a deployment used identical donor subsets, and donor weights were renormalized within each subset.

Let *L*_*r,k*_ denote the Brier loss obtained from ordering *r* at donor count *k*. Improvement relative to the source confidence mapping was defined as

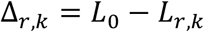

Let Δ_all_ denote the improvement obtained using all target-development donors; this quantity equals the material gap. A donor count met the recovery criterion only when:

1. the 2.5th percentile of Δ_*r,k*_ was greater than 0;
2. the median of Δ_*r,k*_/Δ_all_ was at least 0.25; and
3. at least 80% of valid fits produced *L*_*r,k*_ < *L*_0_.

The minimum donor number was the first tested count that met all three conditions and continued to meet them at every larger tested count. Tested counts were 1, 2, 3, 4, 5, 10, 15, 20 and 30 donors, together with all target-development donors. This value describes the first sustained success on the tested donor grid and is not a universal minimum donor requirement. The same procedure was applied to the Singapore cohort.

### Fixed-cutoff automatic-acceptance audit

To determine how confidence recalibration changes automatic-acceptance decisions, we evaluate three fixed cutoffs:

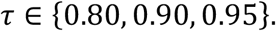

Before recalibration, MSP is used as the decision score. After recalibration, the decision score is the estimated probability of correctness reported by the target confidence mapping. We compare three decision rules:

1. direct use of MSP;
2. a target confidence mapping fitted using five labeled target donors; and
3. a target confidence mapping fitted using all labeled target donors available for recalibration.

Both recalibrated rules use the natural-cubic-spline specification described above. They differ only in the number of labeled target donors used to fit the mapping. Predicted cell types and MSP values remain unchanged.

Let 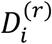 denote the decision score for cell *i* under rule *r*. A prediction is accepted automatically when

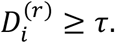

Otherwise, it is referred for manual review. Using reference labels from target donors used only for evaluation, we calculate the number of erroneous automatic acceptances per 1,000 cells as

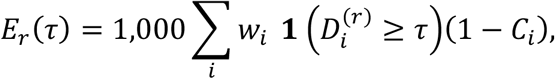

where *C*_*i*_ = 1 for a correct prediction and *C*_*i*_ = 0 otherwise. The corresponding number of manual reviews per 1,000 cells is

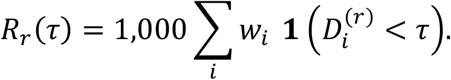

Here, *w*_*i*_ is the normalized donor weight defined above.

For the comparison in Fig. 4B, erroneous automatic acceptances under MSP are set to 100%. The percentage remaining after recalibration is calculated as

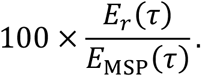

Manual-review burden is reported directly as *R*_*r*_(*τ*). The five-donor rule uses the same 1,000 five-donor samples generated for the finite-donor recovery analysis. For each model–deployment combination and cutoff, we report the median across successful fits. The all-donor rule uses one target confidence mapping fitted with all labeled target donors available for recalibration. The three cutoffs are fixed analysis settings and are not recommended deployment thresholds.

---

**Supplementary Fig. 1.**
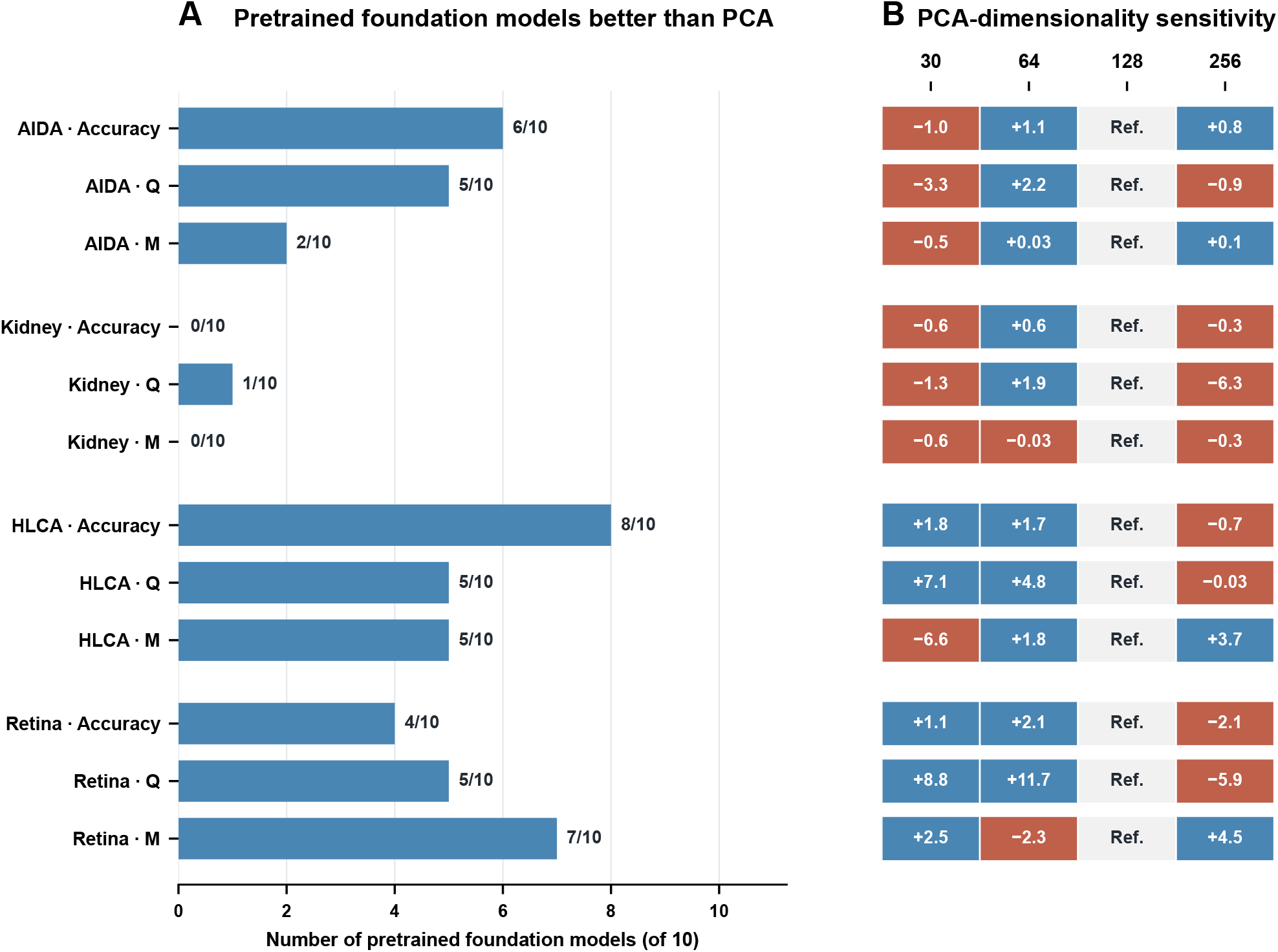
Comparison of single-cell foundation models with PCA and sensitivity to PCA dimensionality. **A**, Within each deployment and metric, bar length indicates the number of pretrained foundation models with point estimates that outperform PCA. Ten models are compared. Higher accuracy indicates better performance, whereas lower *Q* and *M* indicate better performance. **B**, Changes in the metrics for using 30, 64 and 256 principal components (PCs) relative to 128. The three rows within each deployment represent accuracy, *Q* and *M*, respectively. Accuracy differences are reported in percentage points, whereas differences in *Q* and *M* are reported in units of 10^−3^. Positive values indicate that the corresponding PCA dimensionality outperforms 128-dimensional PCA, whereas negative values indicate that 128-dimensional PCA performs better. Blue and red indicate positive and negative values, respectively.

## Notes

### Competing Interest Statement

The authors have declared no competing interest.

